# Molecular detection of virulence and resistance genes in *Salmonella enterica* serovar Typhi and Paratyphi A, B and C isolated from human diarrhea samples and lettuce in Burkina Faso

**DOI:** 10.1101/436501

**Authors:** Namwin Siourimè Somda, Aly Savadogo, Juste Isidore Ouindgueta Bonkoungou, Oumar Traoré, Bissoume Sambe-Ba, Abdoul Aziz Wane, Yves Traoré, Amy Gassama-Sow

## Abstract

**Objectives:** In Burkina Faso (BF), dirty water, in particular those of the stoppings and the gutter are used for irrigation of vegetables. The aim of this study is to contribute to the knowledge on the molecular level of *Salmonella* Typhi and Paratyphi circulating in the hospitals and environment next to hospitals in BF.

**Methods:** *Salmonella* Typhi and Paratyphi strains isolated from patients between 2009 to 2015 and lettuce samples isolated in 2014 in BF were characterized by simple PCR using specific primers.

**Results:** Out of 100 *Salmonella* isolated, 53% were from human and 47% from lettuce samples. Globally, the highest prevalence was observed with *inv*A, *mis*L, *pip*D, *orf*L and *spv*R genes in 97%, 96%; 74%; and 21%. Forty of these isolates carried class 1 integron, 31 from clinical samples and 9 from lettuce samples. Sequencing showed seven different gene cassette arrangements, with *aadA1* in 13/15 strains, *aadA7* and *aac(3)-Id* in 2/15 strains. Eight percent (8/100) of *Salmonella* harbored *gyrB* and *parE* genes with 6 from clinical and 2 from lettuce isolates. Sequencing showed no mutation in these genes. Three distinct PFGE types were observed from clinical samples with 90-95% similarity in each case. All *Salmonella* from lettuce had similar pulsotypes.

**Conclusion:** This study showed the diversity virulence and resistance genes harbored of *S.* Typhi and Paratyphi from both clinical and lettuce samples in BF. Lettuce is a potential source of transmission of *Salmonella* causing diarrhea among human in BF.

## 1. Introduction

*Salmonella entrica* are bacteria that cause salmonellosis and are common causes of human foodborne outbreaks and diseases in developed and developing countries with attendant public health problem [1]. It can be disseminated by industrial water, agricultural, domestic, sweet and drinking, groundwater and seawater. Their transmission to human is principally by ingesting contaminated water or food [2].

In Burkina Faso, rains shortage leads to the practice of farming irrigated by dam or untreated waste water. Untreated dirty water in particular, of the stoppings and the gutter are often used for irrigation of vegetables while these water are nests of pathogenic germs, such as *Salmonella*. These practices lead to the production of vegetables such as lettuce contaminated with pathogens. Consumption of these products results in diarrheal diseases. To treat these infections in developing countries, fluoroquinolones (FQ) and third-generation cephalosporins have been considered as first-line drugs [3]. In recent years, with multidrug resistance, another trend has arisen: the emergence of virulence-resistance (VR) plasmids; these are hybrid plasmids that harbor resistance (R) and virulence (V) determinants [3]. The appearance of these plasmids is of concern because they could lead to the co-selection of virulence (in addition to resistance) through the use of antimicrobial drugs [4]. It could result not only an increase virulence and resistance genes in some strains but also promote the transfer of genes virulence and resistance between strains of the same species or different species. In BF there is a lack of data on the epidemiology of virulence and resistance genes in *Salmonella* Typhi and Paratyphi isolated from diarrheal stools and food. This is the first report of the prevalence of these genes in BF. This study contributes to the knowledge at the molecular level of epidemiology, virulence and resistance genes of *Salmonella* Typhi and Paratyphi circulating in the hospital and environmental circles in BF.

The aims of this study were (i) to characterize the virulence and resistance genes of *Salmonella* Typhi and Paratyphi isolated from stool samples and lettuce in BF, (ii) to determine mechanisms of transfer of resistance genes (conjugation) of the same species or strains of a species to another and (iii) the molecular strain typing.

## 2. Material and methods

### 2.1. Collection of bacterial strains

A total of 47 *Salmonella* Typhi and Paratyphi A, B, C were isolated from lettuce samples collected in 2014 in the surrounding environments of the dam number 3 of Ouagadougou and the university hospital Yalgagado Ouédraogo, the biggest hospital in Burkina Faso. In addition, 53 *Salmonella* Typhi and Paratyphi A, B, C were isolated from children with diarrhea between 2009 and 2015 in three regions of Burkina Faso:

- Ouagadougou (*Hopital du District de Bogodogo* (HDB) and *Laboratoire National de Santé Publique* (LNSP)), which is the capital city located in the center of Burkina Faso;
- Gourcy (*District Sanitaire de Gourcy (DSG*)) and
- Boromo (*District Sanitaire de Boromo (DSB*)), which are rural areas located in northern and western parts of the country.

All *Salmonella* strains were stored at −30°C for molecular biology analysis to *Institut Pasteur of Dakar (IPD)*, Senegal.

### 2.2. Antimicrobial susceptibility test

All strains were subjected to a set of 14 antibiotics to study their antibiogram by using Kirby-Bauer disk diffusion method.

### 2.3. DNA extraction

Three colonies of each isolate on nutrient agar plate were picked and suspended in 200μl of distilled H_2_O. After vortexing, the suspension was boiled for 10 minutes, and 50μl of the supernant was collected after spinning for 10 minutes at 12148 rpm in a microcentrifuge as described by Madadgar et al. (2008) [5].

### 2.4. PCR amplification and sequencing

***Virulence genes detection:*** distribution of five genes and plasmids (*invA, spv*R*, orfL, pipD* and *misL*) associated with the virulence of *Salmonella* was determined. PCRs were carried out with theirs specific primers.

***Integron and Resistance genes detection:*** distribution of integrons and fifteen genes associated with the resistance of quinolones and beta-lactams of *Salmonella* was determined. PCRs were carried out with the specific primers *intI*1*, intI*2 *and intI*3 [7], *gyrA, gyrB, parC, parE, qnrA, qnrB, qnrS, qepA* [6], *bla*CTX-M1, *bla*CTX-M2, *bla*CTX-M9, *bla*TEM, *bla*SHV as desrcribed by Ploy et al. 2000 [7].

PCR was performed in a reaction of 25 μl containing 2.5μl 10 X PCR buffer (SolisBiodyne, 25FMG0560), 0.75μl dNTPs (10mM), 0.75 μl MgCl2 (50 mM), 0.2μl Taq DNA polymerase (5 U/μl) and 25ng of DNA sample. PCR was carried out by a thermocycler (Applied Biosystem, 2720 thermal cycler, Singapore) and amplified products were resolved in a 2% agarose gel and visualized under UV light.

***Sequencing:*** It was performed using the BigDye terminator protocol on the ABI PRISM 310 Sequencing sequencer according to the manufacturer (Perkin-Elmer Applied Biosystems, Les Ulis, France). The sequences obtained corresponding were searched using the BlastN program available in http://www.nbci.nlm.nih.gov.

### 2.5. Bacterial conjugation

A total 21 *Salmonella* Typhi and Paratyphi A, B, C (18 for clinical and 3 for lettuce samples) carrying class 1 integrons and resistant to nalidixic acid were used to bacterial conjugation. The recipients (*E. coli* NalR) were selected on Luria-Bertani (LB) medium containing streptomycin (25μg/ml), nalidixic acid (50μg/ml), trimethoprim (5μg/ml) and ampicillin (100μg/ml). The donors were selected on LB containing only streptomycin and trimethoprim (BioMérieux, France). A total of 5 ml of fresh LB broth was inoculated with either recipient or donor bacteria and incubated for 24 h at 37°C. After, cultures were diluted at 1:50 and incubated at 37° C with strong agitation for 4 hours. Donor and recipient strains were then mixed at the following ratios 1:1; 1:2; 1:10 and incubated at 37°C for 3 hours. Transconjugants were selected on LB agar plates supplemented with streptomycin (25μg/ml), nalidixic acid (50μg/ml), trimethoprim (5μg/ml) and ampicillin (100μg/ml). PCR was performed on transconjugants to determine whether resistance genes were transferred.

### 2.6. Pulsed-Field Gel Electrophoresis (PFGE)

A total 29 *Salmonella* Paratyphi B (21 from clinical and 8 from lettuce) were typed. DNA from all *Salmonella* isolated was prepared in agarose plugs [8]. Bacterial cells were embedded in low-melting-point agarose (Biorad, Marnes-la-coquette) and lysed with lysis buffer containing lysozyme and proteinase K. DNA was digested at 37°C with 10U/μl of the restriction endonuclease Xbal (Biorad, Marnes-la-coquette, France). Digested DNA from *S.* Braenderup H9812 was loaded every five lanes as the molecular marker, as recommended by PulseNet [9]. The digests were run at 6V/cm at 14°C for 16 hours in a 1% agarose gel in 0.5X Tris-Borate EDTA buffer using the Genepath system (Biorad, Marnes-la-coquette, France). The gel was stained with ethidium bromide and photographed on an ultraviolet trans-illuminator (GelDoc, Biorad, France). The restriction endonuclease digests were compared visually and isolates with the same pattern of bands (same number and molecular weight) were considered to be the same strain.

## 3. Results

### 3.1. Detection of virulence genes

Out of 100 *Salmonella* Typhi and Paratyphi isolated, 53% were from human and 47% from lettuce samples. Highest prevalence was observed for *invA*, *misL*, *pipD*, *orfL* and *spvR* genes; observed in 97 %; 96 %; 74 %; and 21 % of samples respectively (Table 1).

**Table 1:**
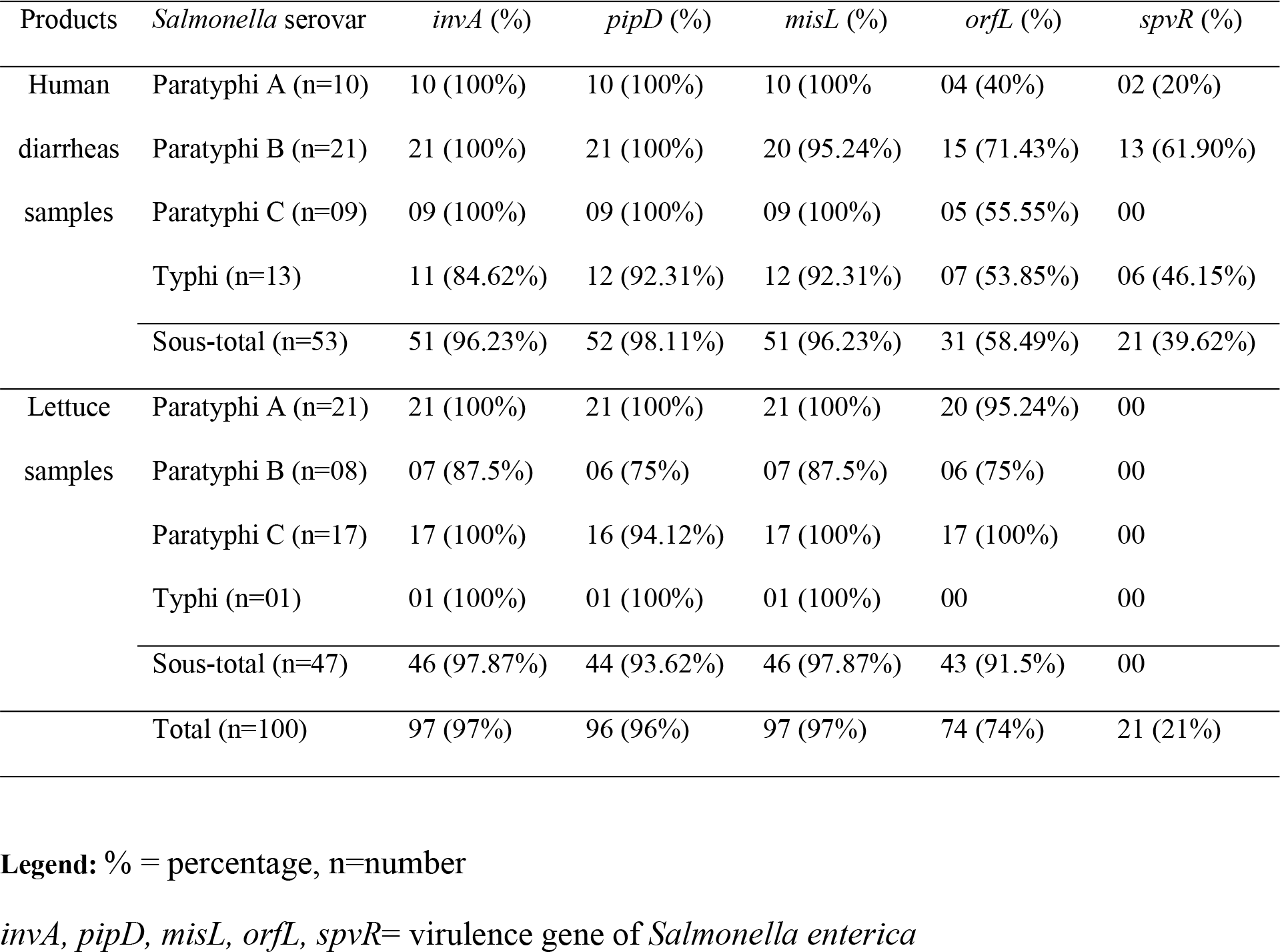
Distrubution of virulence genes of *Salmonella* Typhi and Paratyphi isolated from humans diarrheal samples and lettuce samples in Burkina Faso

### 3.2. Antimicrobial susceptibility test

Among 100 isolates of *S*. Typhi and Paratyphi collected from human and lettuce samples, 40% were resistant at least one antibiotic.

### 3.3. Detection of resistance genes

Forty percent of these isolates carried class 1 integron, with 31 (58.5 %) from clinical and 9 (19.15%) from lettuce samples. Out of these 40 isolates, 15 isolates (14 from clinical and 1 from lettuce sample) had cassettes genes. Sequencing of the variable amplicons of class 1 integron revealed the presence of gene cassettes containing:

- Aminoglycoside acetyltransferase (*aac(3)-Id*) (**GenBank: KT581256.1**) gene which confers resistance against gentamycin,
- Aminoglycoside adenyltransferase (*aadA1, aadA7*) genes (**GenBank: DQ388124.1** and **GenBank: KT581256.1**) that confer resistance to streptomycin and spectinomycin,
- dihydrofolate reductase, (*dfrA1, dfrA7*) (**GenBank: DQ388124.1**and **GenBank: HM769861.1**) that confers resistance to trimethoprim (Table 2). In the variable region of cassettes, different genes were follows organized:

- 1 isolate: *aac(3)-Id* gene followed by *aadA7* gene;
- 13 isolates: *dfrA1* gene followed by *aadA1*gene;
- 1 isolate: *dfrA7* gene followed by 3prime region elements such as *qacE∆1* and *sul1*.

**Table 2:**
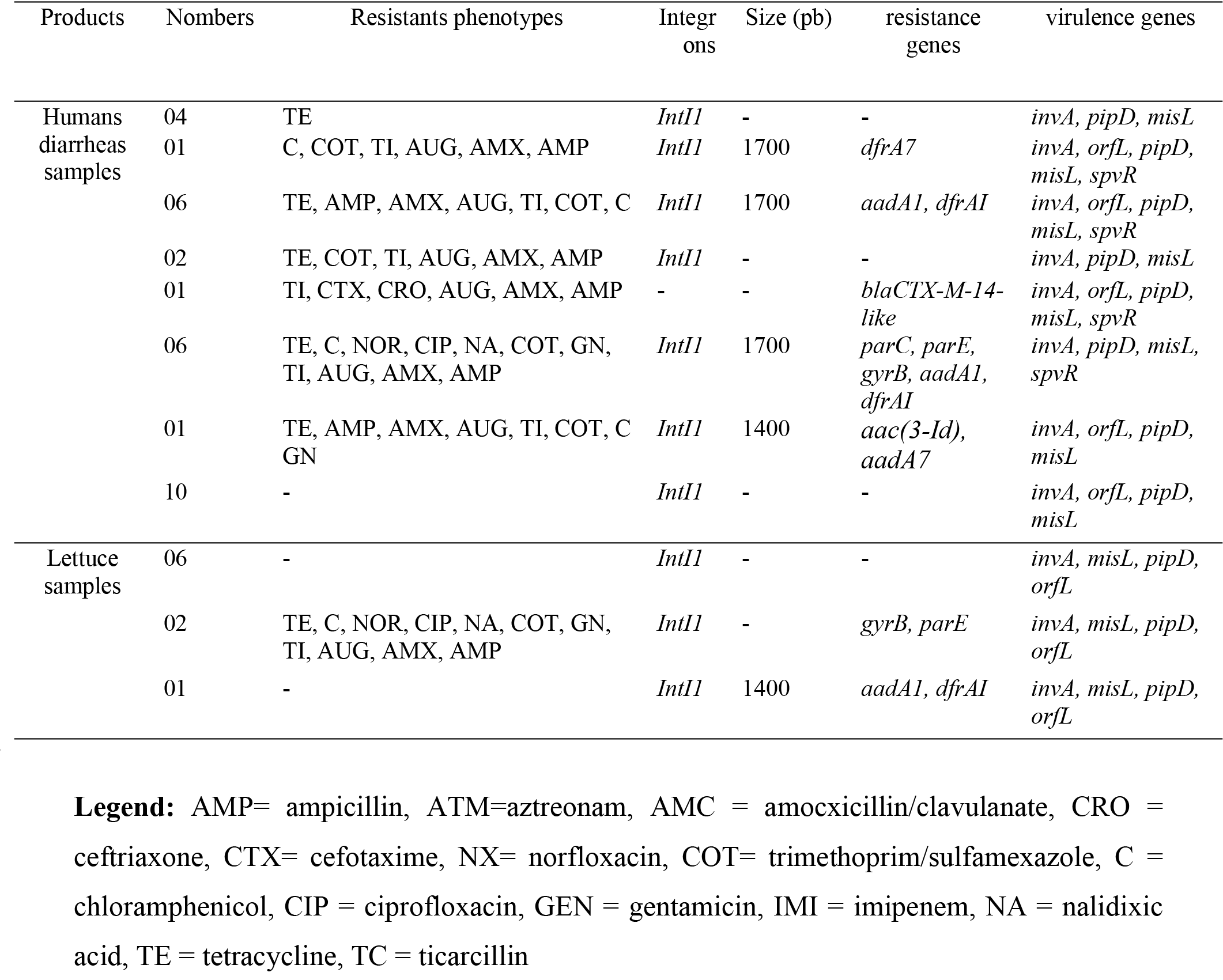
Distrubution of phenotypes resistance, resistance genes and virulence genes of *Salmonella* isolated from human diarrheas and lettuce samples in Burkina Faso.

Interestingly, the *aadA1* and the *dfrA1* were the most predominant resistance genes identified in 13/15 strains carrying class 1 integron cassettes.

Eight percent (8/100) of isolates harbored *gyrB* and *parE* gene (6 from human diarrheas and 2 from lettuce samples) (Table 2). Sequencing of these genes notified no mutation. No *qnr*, *qepA* and *aac(6′)-Ib-c* gene was found in this study. One clinical *Salmonella* Paratyphi B harbored both *aac(3)-Id, aadA7, gyrB*, *parE, parC* genes and blaCTX-M9 gene whose sequencing showed *blaCTX-M-14-like* gene (**GenBank: KX421096.1**). Our study showed that 86.67% (13/15) of strains which harbored class 1 integron genes contain the virulence plasmid *spvR* (Table 2). The majority (9/13) of these strains were *Salmonella* Paratyphi B. The bacterial conjugation showed that resistant genes were not transferable. Transconjugants were absent during this study.

### 3.4. PFGE

The genetic relatedness among the *S.* Paratyphi B strains (n =29 (21 from clinical samples and 08 from lettuce samples)) was investigated by *XbaI*-macrorestriction analysis (PFGE) (Fig. 1). Three distinct PFGE types were observed in each case from clinical samples with 90-95% similarity. All pulsotypes from lettuce sample were similar to 95%. Eighty to ninety (80-90%) similarity was observed with clusters from clinical and lettuce samples (fig. 1). All isolates presenting similar genetic relatedness have similar phenotype resistance and virulence genes.

**Fig 1.**
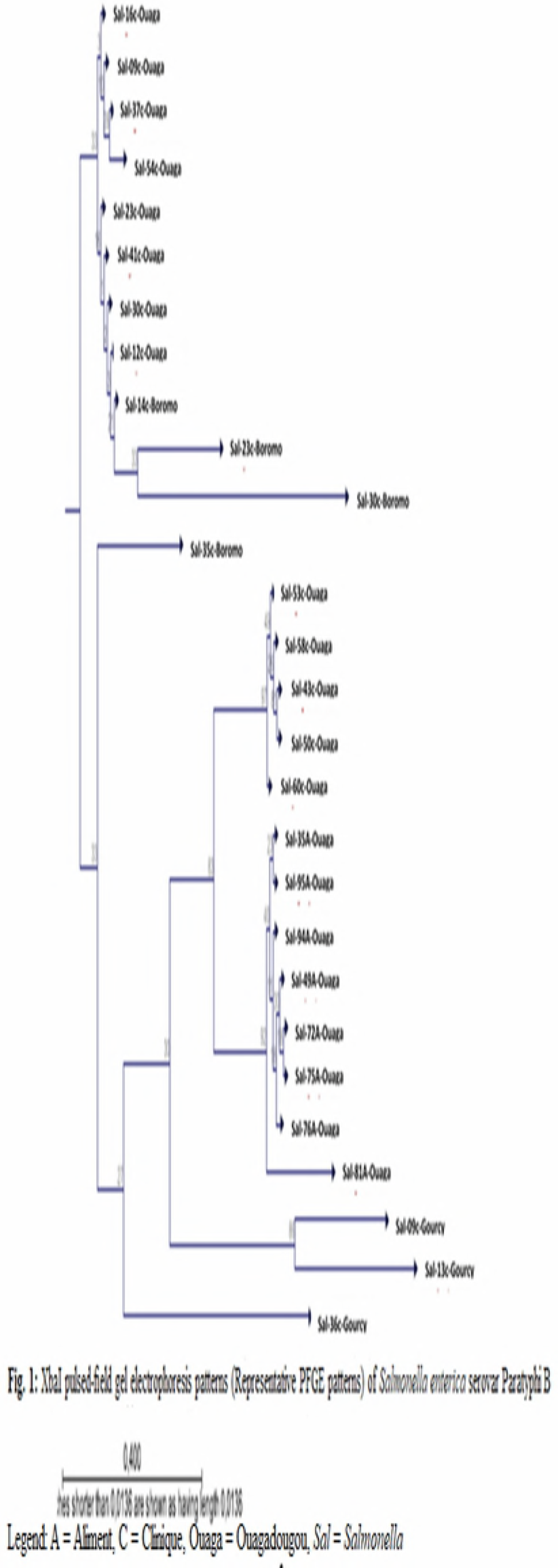
A = aliment, C = Clinique, Ouaga = Ouagadougou, *Sal* = *Salmonella*.

## 4. Discussion

The genome of *Salmonella enterica* possesses multiple pathogenicity islands (PIs), which are genetic elements within the bacterial genome that harbor genes associated with virulence [10]. For our experiments, five virulence genes for PCR amplification from the *Salmonella* serovar were selected. All isolates investigated for the presence of virulence factors revealed the presence of *invA*, *misL*, *orfL*, *pipD*, containing respectively in *SPI-1, SPI-3, SPI-4* and *SPI-5*. To our knowledge, this is the first study reporting the distribution of SPIs in Burkina Faso isolates.

The *invA* genes contain sequences that are unique to *Salmonella* spp. and have been shown to be suitable for specific targets in different diagnostic and research laboratories [11, 12]. Several studies in Africa have shown similar results [13]. In South Africa, Afema *et al*. (2016) found similar results from treated waste water used for the irrigation of vegetables [14].

The *misL* gene located on SPI-3 encodes an auto-transporter protein involved in intestinal colonization and essential for survival in macrophages [15]. The majority of *Salmonella* isolates in this study carried this gene. Similar results were found in most studies in South Africa and Colombia [14, 16].

SPI-4 gene *orfL* was found in 58.49% of human diarrheal strains and in 91.5 % of lettuce strains. SPI-4 is involved in secretion of toxins that induce apoptosis in immune cells and it is required for survival in macrophages while the *pipD* gene encodes effector proteins for the T3SS transport protein and is mainly associated with enteropathogenesis [17]. The macrophages lyse, and bacteria are released to the extracellular space. The significance of this process for *Salmonella* pathogenesis is unknown [15]. Recent studies have shown that caspase-1 apparently has a protective role for the host during systemic *Salmonella* infection, suggesting that caspase-1 activation by *Salmonella* would be detrimental to the organism in disseminated disease [18, 19].

The *spvR* locus is strongly associated with strains that cause non-typhoid bacteremia, but are not present in typhoid strains [15]. *SpvR* is a positive regulator gene of four effector genes, *spvA, spvB, spvC* and *spvD* [15]. These virulence genes are found in the most frequently isolated non-typhoid serotypes of *S. enterica*. Indeed only *Salmonella* Typhimirium and Enteritidis could contain the *spv* plasmids of virulence, which explains why bacteria carrying this plasmids can’t cause gastroenteritis in people [15]. However, this study showed *spvR* in 61.9 % of *Salmonella* Typhi and Paratyphi isolates from human diarrheal samples. The presence of these virulence genes in *Salmonella* Typhi and Paratyphi isolated from lettuce and clinical samples indicate the capabilities of these isolates in causing infections in susceptible hosts.

Resistances related to class 1 integrons were found in 40%. Class 2 and 3 integrons were absents. This predominance of class 1 integrons in *Salmonella* was previously described by other authors [20–23]. According to Vo *et al.* (2010) [24] the prevalence of integrons found in *Salmonella* varies from country to another country and depends on the origin of the isolates. Class 1 integrons were found to 35%; 28%; 46%; 56.72% in Vietnam, England, Kenya and Brazil respectively [24–26]. Sequencing of the gene cassette showed that most of our strain carried *aadA1* and *dfrAI*. Others studies realized in Africa (Senegal, Kenya, Egypt) were showed similar results [23, 25–27]. Resistance to trimethoprim/sulfamexazole is strongly associated with the presence of class 1 integrons, due to the frequent presence of a *dfrA1* cassette and *sul1* gene in the 3’ region [28].

Integrons are potentially capable to transmit drug resistance to other *S. enterica* isolates or to other bacteria. Since integron represents the main vehicle of antibiotic resistance, their presence in *S.* Typhi indicates uninterrupted transfer of drug resistance genes from one organism to another irrespective of their species which are worrisome with respect to the spread of AMR [29].

Class 1 integrons are increasingly described in environmental and animal bacteria. The release of hospital and municipal effluents is the main way to integrate integrons into the environment [25]. In this study according to our methodology, transfer experiments were not successful. It is possible that these class 1 integron which carried resistance genes contained in *Salmonella* isolated were not a transferable elements.

The main mechanism of resistance to quinolones is linked to chromosomal mutations especially in the *gyrA* or *parC* genes and more rarely at the *gyrB* and *parE* genes [30, 31]. This study showed only 8 isolates could be amplified both the *gyrB* and *parE* genes. Previous studies from Brazil and Senegal were showed similar results with our results [30–33].

Extrachromosomal genes *qnr*, *aac(6’)-Ib-cr* and *qepA* were discovered since 2002 and are carried by conjugative plasmids [34]. Since then, several types of *qnr* (*qnrA, qnrB, qnrS*) have been described [34]. These genes were not detected in this study. Several studies shown that these plasmids were most found in non-typhoidal *Salmonella* and absent in *Salmonella* Typhi. This was similar with ours results. Others studies shown that plasmids carrying genes encoding ESBLs generally carry genes encoding quinolone resistance [35, 36]. This study showed only one ESBL-producing strain (*blaCTX-M-14-like*). This results were similar to Al-Emran *et al.* (2016) [37] in Sub-Saharan African from blood samples to febrile patients.

Pulsed-field gel electrophoresis has been widely used to determine strain relatedness, to confirm bacterial disease outbreaks, and to identify the sources of strains [38, 39]. In the present study, three distinct PFGE types (pulsotypes) were observed from clinical samples with 90-95% similarity in each case. The diversity of pulsotypes could explained by the fact that the samples come from different zones and the fact that the clinical isolates were collected from much longer time span, so the PFGE patterns would change more. These zones differ from their climatic, socio-cultural and even demographic conditions. However all *Salmonella* Paratyphi B from lettuce samples indicated indistinguishable pulsotypes. This is not surprising since these isolates originate from the same sampling site. In addition, 80-90% similarity was observed between these pulsotypes and clinical ones from Ouagadougou. This could be due to the fact that our strains from lettuce were collected in gardens close to a health center.

## Conclusion

This study showed that the isolated *Salmonella* strains had several virulence factors that play a role at different stages of the infectious process. The study showed the emergence of *Salmonella* resistant to fluoroquinolone which was previously considered as a hospital problem. These pathogens affect humans through contaminated water and/or food. The foods like lettuce might the route of transmissions of *Salmonella* disease in Burkina. It is necessary to have a transversal monitoring by pooling the efforts of the actors of the environment, human and veterinary medicine.

## Acknowledgements

The authors gratefully acknowledge the “LNSP/BF”, “UBE” in Institut Pasteur of Dakar/Senegal and the “LaBIA/UFR/SVT, Université Ouaga I Pr Joseph KI-ZERBO, BF”.

## Authors’ contributions

SNS carried out the sampling and strains isolation, serotyping and their virulence and resistance gene characterize and drafted the manuscript, SA, BOJ, TO, BSB, WAA, TY and GSA supervised the sampling and strains isolation, serotyping, antibiotics susceptibility and participated in writing the manuscript. All authors read and approved the final version of the manuscript.

## Accession Numbers

- *aac(3)-Id* gene: **GenBank: KT581256.1**
- *aadA1* gene: **GenBank: DQ388124.1**
- *aadA7* gene: **GenBank: KT581256.1**
- *dfrA1* gene: **GenBank: DQ388124.1**
- *dfrA7* gene: **GenBank: HM769861.1**
- *blaCTX-M-14-like* gene: **GenBank: KX421096.1**

